# Imposing a curfew on the use of screen electronic devices improves sleep and daytime vigilance in adolescents

**DOI:** 10.1101/259309

**Authors:** Aurore A. Perrault, Laurence Bayer, Matthias Peuvrier, Alia Afyouni, Paolo Ghisletta, Celine Brockmann, Mona Spiridon, Sophie Hulo Vesely, Stephen Perrig, Sophie Schwartz, Virginie Sterpenich

## Abstract

The use of screen electronic devices (SED) in the evening negatively affects sleep. Yet, sleep is known to be essential for brain maturation and a key factor for good academic performance, and thus is particularly critical during childhood and adolescence. While previous studies reported correlations between SED use and sleep impairments, the causal relationship between SED use and sleep in adolescents remains unclear. Using actigraphy and daily questionnaires in a large sample of students (12 to 19 years old), we assessed SED use and sleep habits over one month, including a two-week baseline phase and a two-week interventional phase, where participants were asked to stop screen use after 9 pm during pre-school nights. During the interventional phase, we found that reduction in time spent on SED after 9 pm correlated with earlier sleep onset time and increased total sleep duration. The latter led to improved daytime vigilance. We also observed that the beneficial impact of the intervention on sleep was influenced by catechol-O-methyltransferase gene (*COMT*) Val158Met polymorphism, which is implicated in the dopaminergic modulation of human behaviors, including wake and sleep regulation. These findings provide evidence that restricting SED use in the evening represents a valid and promising approach for improving sleep duration in adolescents, with potential implications for daytime functioning and health.

**STATEMENT OF SIGNIFICANCE:** With the emergence of smartphones and other connected devices, adolescents spend a lot of time on screen electronic devices (SED), especially during the evening. We report that time spent on SED after 9 pm negatively correlates with sleep onset time, sleep duration as well as mood, body weight, and academic performance. Such observable correlations urge for educational strategies to address the chronic lack of sleep observed in today’s adolescent populations. Here we also show that limiting the use of SED after 9 pm improves sleep duration and daytime vigilance in most adolescents. This simple education recommendation pertaining to sleep hygiene can be implemented by every household, yielding direct positive effects on sleep, and presumed benefits for health and daytime functioning.

## INTRODUCTION

In recent years, there has been a spectacular expansion in the use of screen electronic devices (SED), especially in “Digital Natives” teenagers who represent the first generation to be born in the digitalized world and have lived their entire teenage years with access to SED (1). With the proliferation of different types of SED (e.g., laptops, smartphones, tablets) and the diversity of activities that they offer (from blogs and social media to video games), adolescents are “over-connected” (2, 3). Excessive time spent on screen-based activities has not only been considered as one form of technological addiction (4, 5), but has also been shown to affect academic outcome (2) and increase the risk of developing health problems, such as obesity, insomnia or depression (6, 7).

Sleep habits during adulthood and adolescence have also changed in the past years (8). Several studies have reported later bedtime, shortened sleep duration, and longer sleep onset latency (9, 10). In their large sample of Australian teenagers (*N*=1287), King and colleagues (2014) found that most participants reported sleep disturbances as a consequence of SED use, with bedtime delay being the most prevalent problem (11). Other similar studies using questionnaires revealed a negative relationship between time spent on SED and sleep duration (8, 12). Cain and Gradisar (2010) have suggested three possible mechanisms that may contribute to the influence of SED on sleep: i) screen-based activities are time-consuming, and thus compete for time for evening sleep (11, 14); ii) screen-based activities can increase emotional arousal prior sleep, impacting bedtime hour but also sleep onset latency (15, 16); iii) the light emitted by the screens may be interfering with sleep by delaying hormonal melatonin production (17–19). The delay in sleep onset will consequently shorten sleep duration, as wake up time remains unchanged due to fixed school hours. The negative impact of SED use on sleep quantity and quality during childhood and adolescence may have detrimental consequences on their future adult life. Indeed, insufficient and disturbed sleep at a young age is associated with a greater risk to develop obesity, hypertension and mood disturbances, including depression (20, 21). In addition, it is well known that chronic sleep restriction directly affects daytime functioning including vigilance, learning, and executive functions (22), which, during development, may affect performance at school. Actually, sufficient and good sleep have been defined as key contributors to good academic performance (23– 25). Hence, with the continuous expansion of SED and their increasing use, especially prior to sleep, it is critical and urgent to find targeted preventive measures to preserve good sleep.

In the present prospective interventional study performed on a large sample of adolescents, we tested whether imposing a restrictive use of SED after 9 pm beneficially affects sleep. We first wanted to confirm the relationship between SED use in the evening and sleep parameters, using objective (i.e., actigraphy, melatonin profile) and subjective (i.e., questionnaires, self-report diaries) measures. Second, we hypothesized that a reduction of SED use after 9 pm would advance bedtime hour, which in turn should improve sleep duration and daytime functioning. Third, we investigated whether individual catechol-O-methyltransferase (*COMT*) Val158Met polymorphism might modulate the impact of a restrictive use of SED on sleep. *COMT* gene has an established role in cognition and in the maintenance of behavioral arousal through its influence on dopamine activity in the prefrontal cortex (26). *COMT* polymorphisms contribute to variabilities across diverse dopaminergic functions including motivation for reward (27), executive functions (28) and emotion processing (29), with clinical implications such as risk for schizophrenia (30), bipolar disorder (31), or addictive behaviors (32). Although its implication in sleep-wake regulation is not yet fully understood (26, 33, 34), *COMT* polymorphism was shown to predict individual differences in sleep changes resulting from chronic sleep restriction (35), and to underlie inter-individual differences in brain oscillation during wake and sleep states (36, 37). For all these reasons, we tested whether *COMT* polymorphism influences the relation between SED use and sleep parameters.

## RESULTS

We recruited 569 adolescents who participated in a baseline phase during two weeks (i.e., no change in SED use; Phase 1) and then a second equivalent period during which participants were asked to refrain from using SED after 9 pm (Phase 2; **Fig. 1**). We collected sleep data and time spent on evening activities on- and off-screen using daily questionnaires and actigraphy. Vigilance measurement and saliva samples for melatonin profiling and genetic analysis (COMT) were also obtained at baseline and after the intervention. Data analyses were performed on “Active” participants (i.e., those who completed questionnaires, wore actimeter, and filled out daily diaries for at least 7 days). We called participants who did not fulfil these criteria “Passive” and only analyzed their questionnaire and vigilance data.

**Figure 1:**
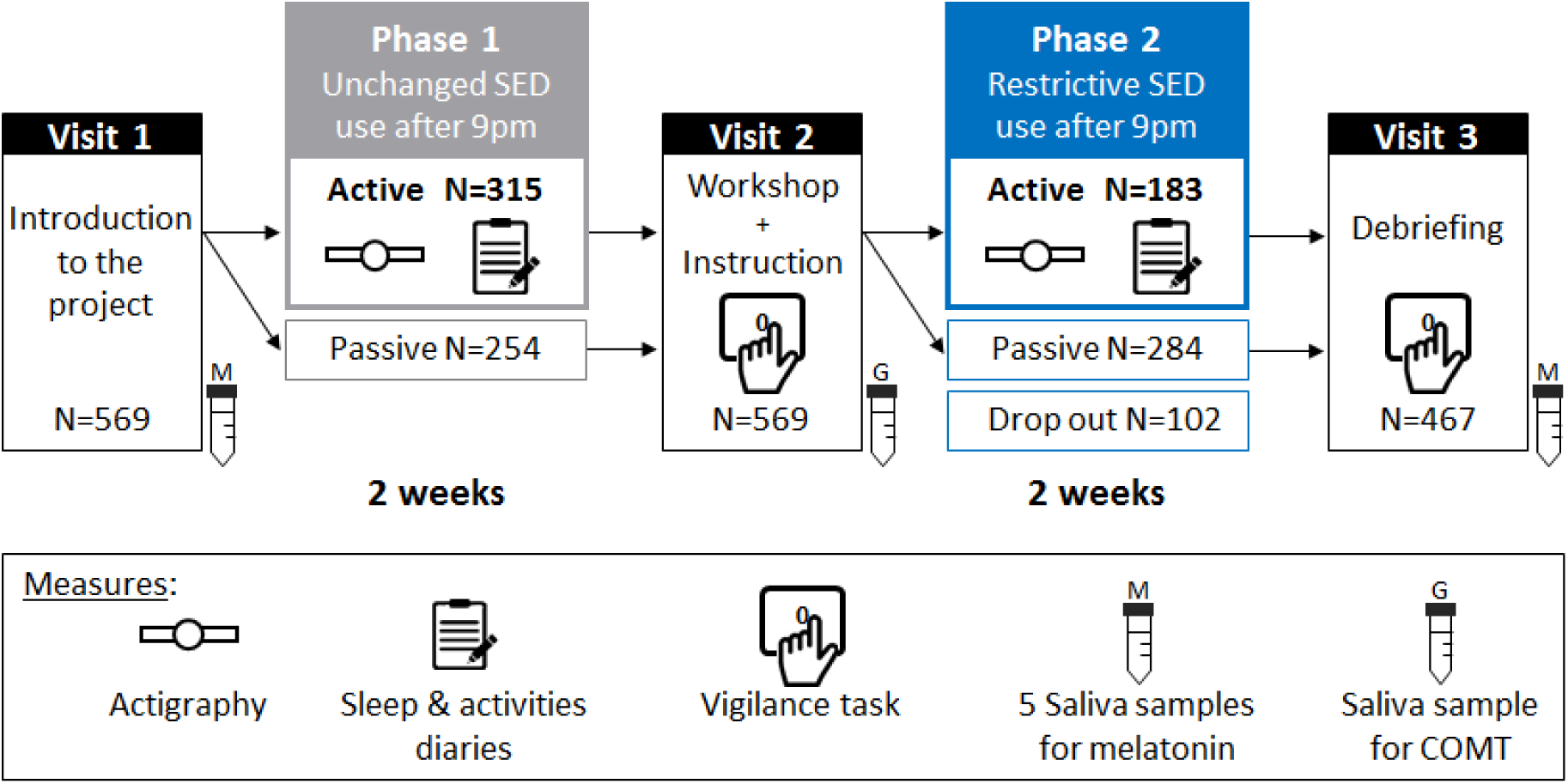
Study design and participants. Repartition of the 569 adolescents across the protocol: 2×2 weeks, including baseline period (no change in SED use; Phase 1) and experimental period (restricted use of SED after 9 pm; Phase 2), where we collected sleep (actigraphy) and evening activities (diaries) data. Vigilance and saliva samples (for melatonin and genetic profiles) were also obtained at baseline and after the intervention.

### Phase 1

#### On- and off-screen activities after 9 pm

The intervention explicitly targeted screen-based activities during the evenings preceding school days (i.e., Sundays to Thursdays; see Methods). We therefore first focused on the pre-school evenings and nights measurements. During Phase 1, 96.8 % of the *Active* participants (*N*=315) reported spending on average (± SEM) 79 (± 3) min on SED after 9 pm on pre-school evenings. ANOVA on time spent on different Types of SED (Social media, TV, Videos, Games, Computers) as within-subject factor and Age Group (12-13, 14-15, 16-17, 18-19 years old) as between-subject factor revealed a main effect of Types of SED (*F*(4,1244) = 40.46; *P* <.0001) because they exhibited higher preference for social media, on which they usually spent more than 18% of their time between 9 pm and sleep onset. Moreover, there is a main effect of Age Group (*F*(3,311) = 12.81; *P* <.0001) as older teenagers spent more time on SED than younger ones. Finally, the interaction between SED type and age was also significant (*F*(12,1244) = 2.41; *P* =.004) because the use of different types of SED evolved disparately with age (**Fig. 2*A***). Similar ANOVA on time spent on off-screen activities (Homework, Reading, Sport) revealed a main effect of Types of Off-screen (*F*(2,622) = 403.4; *P* <.0001) due to larger amount of time spent doing homework. However, we observed no main effect of Age Group (*F*(3,311) = 2.23; *P* =.084) but a significant interaction (*F*(6,622) = 2.7; *P* =.013) mediated by larger homework duration in older adolescents (18-19 years old; **Fig. 2*B***).

**Figure 2:**
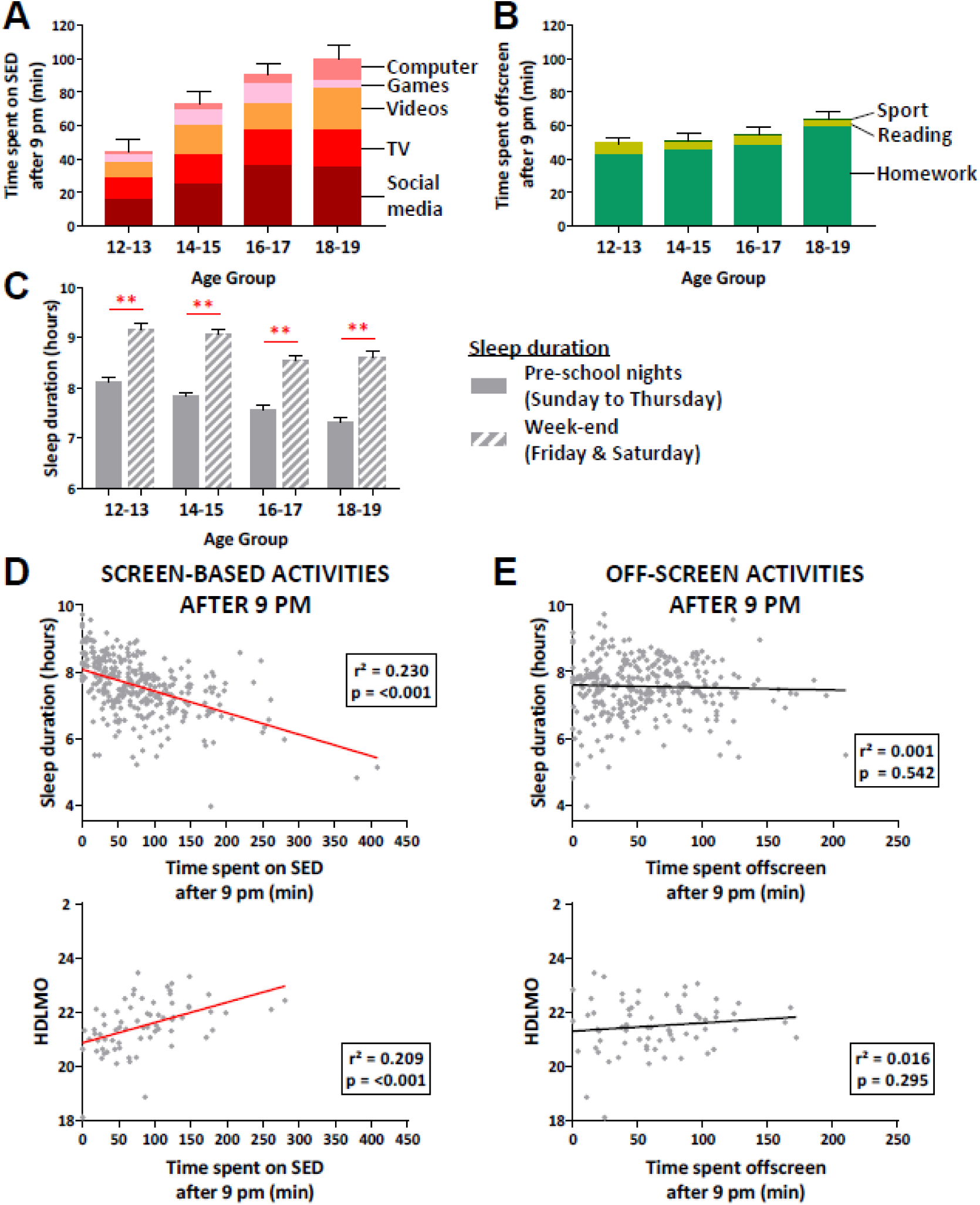
Phase 1-Activities after 9 pm and sleep during pre-school nights. (A) Mean time spent on each screen-based activity (± SEM of total time spent on SED) after 9 pm during pre-school nights per age group. (B) Mean time spent on each off-screen activity (± SEM of total time spent off-screen) after 9 pm during pre-school nights per age group. (C) Mean (± SEM) sleep duration during pre-school nights (grey) and weekend nights (grey strips) per age group. (D) Scatter plots showing significant correlation (*P* <.001) between the time spent on screen-based activities after 9 pm and sleep duration (*N*=315; top), and hour of dim light melatonin onset (HDLMO; *N*=70; bottom) during pre-school nights. (E) Scatter plots showing no correlation (*P* >.5) between the time spent on off-screen activities after 9 pm and sleep duration (*N*=315; top), and HDLMO (*N*=70; bottom) during pre-school nights. Asterisks represent significance (*P*) of 2-tailed paired *t*-tests between pre-school nights and weekend nights: ** <.001

#### Sleep parameters

During Phase 1, sleep duration (or total sleep period; time between sleep onset and wake-up time) for pre-school nights decreased with age (ANOVA on sleep duration with Age Group as between-subjects factor; *F*(3,311) = 27.05; *P* <.0001; **Fig. 2*C***), while sleep onset time was progressively delayed with age (ANOVA on sleep onset with Age Group as between-subjects factor; *F*(3,311) = 19.84; *P* <.0001). On average (± SEM), adolescents slept 7h33 (± 3 min) during pre-school nights. For weekend nights, we observed that adolescents slept longer (8h40 ± 4 min) suggesting a possible sleep debt accumulated during the week. Thus, we performed an ANOVA on sleep duration with Type of Night (pre-school nights, weekend nights) as within-subjects factor and Age Group as between-subjects factor and we observed a significant main effect of Type of Night (*F*(1,298) = 305.59; *P* <.0001). This reflects an extended period of sleep during the weekends, a main effect of Age Group (*F*(3,298) = 24.9; *P* <.0001) as sleep duration differs between age group, although sleep rebound was present across all age groups (all *t*-test *P <.001)*. There was also a significant interaction (*F*(3,298) = 2.65; *P* =.049) due to shorter difference between pre-school nights and weekend night in older adolescents (16-19 years old) compared to younger adolescents (**Fig. 2*C***).

#### Link between evening activities, sleep and waking performance

We next tested whether activities performed during pre-school evenings influenced subsequent sleep. We first found that total time spent on SED after 9 pm correlated negatively with sleep duration (*R*^*2*^ = 0.23; *P* <.001; **Fig. 2*D***). By contrast, total time spent doing off-screen activities did not significantly affect sleep duration (*R*^*2*^ = 0.001; *P* =.54; **Fig. 2*E***). Note that because wake-up time is constrained by early morning school schedules, the impact of SED use on sleep duration was primarily attributable to a later sleep onset time. Indeed, there was a significant correlation between the time spent on SED after 9pm and sleep onset time (*R*^*2*^ = 0.35; *P* <.001). Finally, there was a significant correlation between the duration of screen-based activities in the evening and melatonin profiles (Hour of Dim Light Melatonin Onset, HDLMO; see Methods; *R*^*2*^ = 0.239; *P* =.001; **Fig. 2*DE***) suggesting a direct impact of exposure to SED light on the circadian regulation of sleep. Note that similar correlations during weekend nights did not reveal significant correlation between total time spent on SED after 9 pm and sleep duration (*R*^*2*^ = 0.002; *P =*.35). To better understand how different types of media might influence sleep duration across the distinct age groups, we used a Structural Equation Modeling (SEM) approach, which revealed that social media, videos, games, and computer activities significantly affected sleep duration (all *P* <.001). Watching TV (modulated by age) was the only media that did not affect sleep duration (*R*^2^** = 0.086, *P* =.11). Computer use seemed to have the highest unique impact on sleep duration as it explained 6.3 % of the variance (compared to 4.8 % for video games; 2.4 % for watching videos, and 1.5 % for social media; **Fig. S1**). Note that the model obtained a good fit (χ^2^ =13.32, df =10, *P* =.2, RMSEA =.033, CFI =.985).

Regarding waking performance, extensive SED use in the evening correlated with lower grades at school (*R*^2^** = 0.064; *P* <.001), increased psychological distress (K6; *R*^2^** = 0.014; *P* =.033), increased daytime fatigue (CSRQ; *R*^2^** = 0.033; *P* =.001), and higher BMI score (*R*^2^** = 0.046; *P* <.001). However, no correlation was observed between SED use and performance on the SART (Sustained Attention to Response Task), an objective measure of vigilance (*P* >.05 for all SART measures; see Methods). Note that sleep duration (which was influenced by SED use, see above) did not correlate with school performance, nor psychological distress, but correlated negatively with daytime fatigue (CSRQ; *R*^2^** = 0.065; *P* <.001), BMI score (*R*^2^** = 0.073; *P* <.001), and daily mood rating (*R*^2^** = 0.016; *P* =.024).

### Comparisons between Phase 1 and Phase 2

#### Effects of the intervention on on- and off-screen activities after 9 pm

On average, *Active* participants (*N*=183) reduced their time spent on SED by 71.3 % after 9 pm on pre-school evenings (mean ± SEM; Phase 1: 76.15 ± 3.57 min, Phase 2: 21.49 ± 2.15 min; **Fig. 3*A***). An ANOVA on SED use duration after 9 pm using Phase (Phase 1, Phase 2) as within-subjects factor and Age Group as a between-subjects factor revealed a significant main effect of Phase (*F*(1,179) = 220.73; *P* <.0001) due to the reduction in SED use during Phase 2. This effect is present in each age group (post-hoc *t-*test, all *P* <.001). We observed a main effect of Age Group (*F*(3,179) = 7.95; *P* <.0001) and a significant interaction (*F*(3,179) = 12.44; *P* <.001) as older adolescents (14-19 years old) exhibited larger use of SED during baseline and therefore exhibited greater reduction during Phase 2 (**Fig. 3*B***). A second ANOVA on Phase and Types of SED similarly revealed a main effect of Phase (*F*(1,182) = 190.1; *P* <.0001) and Types of SED (*F*(4,728) = 34.5; *P* <.0001) as all types of screen-based activities were reduced. There was also a significant interaction (*F*(4,728) = 25.8; *P* <.0001) explained by the larger reduction in social media, watching TV and videos which were the most used activities during baseline.We used similar analyses to check the impact of the intervention on off-screen activities after 9 pm. ANOVA using Phase and Age Group showed no main effect of Phase (*F*(1,179) = 1.23; *P* =.26), a main effect of Age Group (*F*(3,179) = 3.08; *P* =.028), but no interaction (*F*(3,179) = 0.85; *P* =.46) suggesting stable time dedicated to off-screen activities between phases for each age group. Moreover, when looking at the different types of off-screen activities, we found a main effect of Types of Off-screen (*F*(2,364) = 207.4; *P* <.001) due to large amount of time doing homework in both phases and a Phase by Types of Off-screen interaction (*F*(2,364) = 19.9; *P* <.001) that was driven by a significant increase in the time spent reading (mean ± SEM; Phase 1: 5.43 ± 0.45 min, Phase 2: 14.18 ± 1.30 min; *t*-test P <.001) during Phase 2.

**Figure 3:**
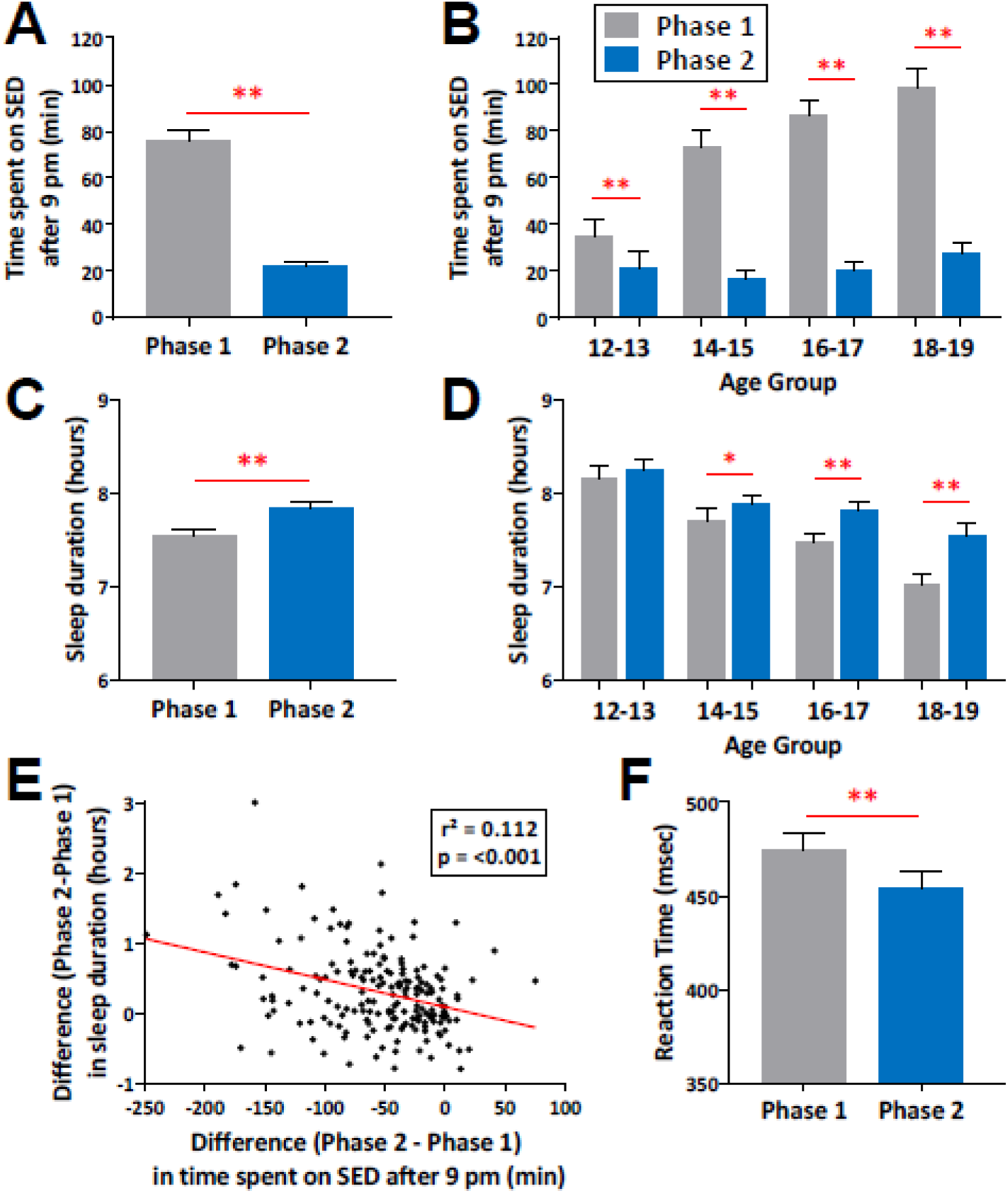
Phase 1 and 2 - Impact of the restrictive use of SED after 9 pm on sleep and vigilance. (A) Mean (± SEM) time spent on screen-based activities after 9 pm during pre-school nights for Phase 1 (grey) and Phase 2 (blue). (B) Mean (± SEM) time spent on screen-based activities after 9 pm per age group for Phase 1 and Phase 2. (C) Mean (± SEM) sleep duration during pre-school nights for Phase 1 and Phase 2. (D) Mean (± SEM) sleep duration per age group for Phase 1 and Phase 2. (E) Scatter plot showing a significant correlation (*P* <.001) between the difference in time spent on screen-based activities after 9 pm (Phase 2 minus Phase 1) and the difference in sleep duration (Phase 2 minus Phase1) during pre-school nights. (F) Mean (± SEM) reaction time (slowest 75^th^ percentile) during the vigilance task (SART) performed at the end of Phase 1 (grey) and Phase 2 (blue). Asterisks represent significance (*P*) of 2-tailed paired *t*-tests between Phase 1 and Phase 2: ** <.001, *<.05

#### The restrictive use of SED beneficially impacts sleep parameters

*Active* participants (*N*=183) during Phase 1 and 2 also went to bed earlier on pre-school nights (mean ± SEM; Phase 1: 23h28 ± 4 min, Phase 2: 23h07 ± 3 min; *t-*test *P* <.001) and consequently increased their sleep duration (mean ± SEM; Phase 1: 7h33 ± 3 min, Phase 2: 7h50 ± 3 min; *t-*test *P* <.001; **Fig. 3*C***) during Phase 2. An ANOVA on sleep duration using Phase as within-subjects factor and Age Group as a between-subjects factor revealed a significant main effect of Phase (*F*(1,179) = 44.03; *P* <.001), Age Group (*F*(3,179) = 11.14; *P* <.001) and a Phase by Age Group interaction (*F*(3,179) = 5.23; *P* =.002) due to significant increase in sleep duration between phases for adolescents between 14 and 19 years old (*P* <.05; **Fig. 3*D***). Note that same ANOVA on sleep onset hour revealed similar significant results (all *P* <.0001) demonstrating earlier bedtime during Phase 2. The suggested relationship between decreased SED use in the evening and subsequent sleep was further supported by significant correlations between the difference (between Phases 1 and 2) in time spent on SED and the difference in sleep onset time (*R*^2^** = 0.133; *P* <.001), and the difference in sleep duration (*R*^2^** = 0.112; *P* <.001**; Fig. 3*E***). In summary, the better participants complied with the instructions regarding SED use, the earlier they went to bed and the more they slept. There was no significant change in melatonin profiles between Phases (*t*-test *P* =.59). Note however that we could obtain reliable melatonin profiles for both Phase 1 and Phase 2 from only 13 participants.

#### Restrictive use of SED during the evening affects daytime vigilance

*Active* participants of Phase 2 who performed the SART task during Visit 2 and Visit 3 (*N*=177; **Fig. 1**) exhibited faster reaction times on slowest RTs (75^th^ percentile) in Phase 2 compared to Phase 1 (mean ± SEM; Phase 1: 474.21 ± 9.31 ms, Phase 2: 453.92 ± 9.6 ms; *P* =.003; **Fig.3*F***). To account for possible learning effects, we compared the results from those participants to the *Passive* participants who also performed the SART in both Phases (*N*=253) with an ANOVA on slowest RTs, using Phase as within-subjects factor and Participation Group (*Active, Passive*) as a between-subjects factor, and observed a significant main effect of Phase (*F*(1,444) = 8.75; *P* =.003) but no effect of Group (*F*(1,444) = .042; *P* =.83) nor Phase by Participation Group interaction (*F*(1,444) = .82; *P* =.36). Critically, note that a *t*-test comparing Phases revealed no significant improvement in the slowest RTs in the *Passive* group (P =.13). Regarding others’ waking performance, we found no significant impact of the intervention on the participants’ self-reported daily mood between Phase 1 and Phase 2 (*t*-test *P* =.094). However, we found a decrease in daytime fatigue (CSRQ) score between phases in both *Active* and *Passive* participants. ANOVA using Phase as within-subjects factor and Participation Group as between-subjects factor revealed a main effect of Phase (*F*(1,444) = 56.42; *P* <.001), no effect of Participation Group (*F*(1,443) = 0.91; *P* =.34) and a Phase by Participation Group interaction (*F*(1,1) = 4.86; *P* =.028) due to larger decrease in CSRQ score, reflecting less fatigue, in *Active* participants after Phase 2.

#### COMT gene polymorphism influences the impact of the intervention on sleep parameters

After genotyping the A/G SNP in exon 3 of *COMT* (rs 4680) using the TaqMan allelic discrimination method (See Methods), we obtained the following distribution: 37 Val/Val homozygotes, 61 Val/Met heterozygotes and 23 Met/Met homozygotes. ANOVAs with *COMT* polymorphisms (Val/Val, Val/Met, Met/Met) as between-subjects factor revealed no significant difference for age, gender, BMI, daily mood rating, sleep parameters, and SED use after 9 pm during pre-school nights during Phase 1 (all *P* >.05; **Table S1**). An ANOVA on time spent on SED after 9 pm with Phase as within-subjects factor and *COMT* polymorphisms as between-subjects factor showed a main effect of Phase (*F*(1,118) = 147.8; *P* <.001) but no effect of *COMT* (*F*(2,118) = 2.7; *P* =.07) nor significant interaction (*F*(2,118) = 0.27; *P* =.76), thus suggesting that all participants decreased their SED use during Phase 2, irrespective of *COMT* polymorphism (**Fig. 4*A***). A similar ANOVA on sleep onset time revealed a main effect of Phase (*F*(1,118) = 40.3; *P* <.001), no effect of *COMT* (*F*(2,118) = 1.003; *P* =.37) but an interaction Phase by *COMT* polymorphism (*F*(2,118) = 5.55; *P* =.005). *Post-hoc t*-tests revealed that both Val/Val (mean ± SEM; Phase1: 23h36 ± 8 min, Phase 2: 23h07 ± 7 min; *P* <.001) and Val/Met (mean ± SEM; Phase 1: 23h25 ± 6 min, Phase 2: 23h03 ± 6 min; *P* <.001) significantly advanced their sleep onset time between Phase 1 and Phase 2, whereas Met/Met carrier did not (mean ± SEM; Phase1: 23h30 ± 7 min, Phase 2: 23h27 ± 8 min; *P* =.54; **Fig. 4*B***). Based on this result, we could expect an impact on sleep duration but the ANOVA on sleep duration revealed only a main effect of Phase (*F*(1,118) = 26.84; *P* <.001) and no effect of *COMT* (*F*(2,118) =.54; *P* =.58) nor interaction (*F*(2,118) = 1.14; *P =*.32). Although the interaction was not significant, we performed exploratory *t*-tests between phases for each *COMT* polymorphism separately and found that, similar to sleep onset time, sleep duration significantly increased between Phase 1 and Phase 2 both for Val/Val (Phase 1: 7h25 ± 9min; Phase 2: 7h46 ± 7min; *P* <.001) and Val/Met (Phase 1: 7h34 ± 6min; Phase 2: 7h52 ± 5min; *P* <.001) adolescents. However, for Met/Met teenagers, despite a decreased SED use, sleep duration was not increased (Phase 1: 7h28 ± 9min; Phase 2: 7h37 ± 9min; *P* =.143).

**Figure 4:**
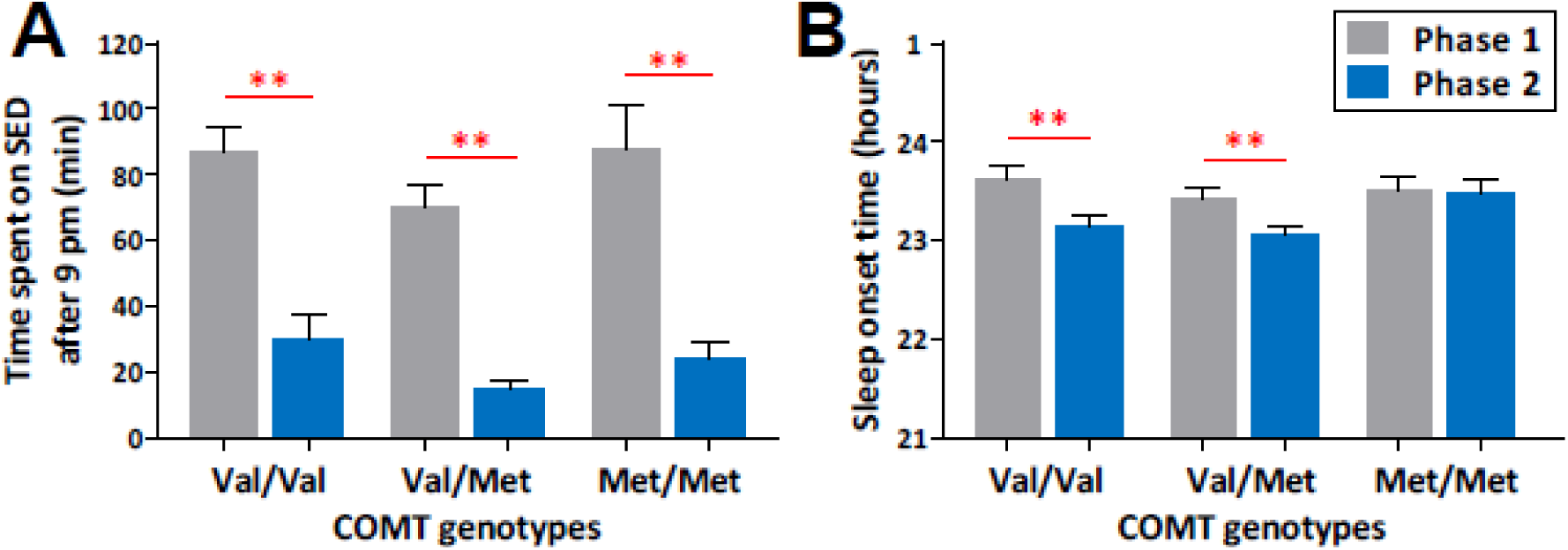
Phase 1 and 2 - Impact of limiting SED use after 9 pm on sleep across distinct COMT polymorphisms. (A) Mean (± SEM) time spent on screen-based activities after 9 pm during pre-school nights per COMT genotype for Phase 1 (grey) and Phase 2 (blue). (B) Mean (± SEM) sleep onset time during pre-school nights per COMT genotype for Phase 1 and Phase 2. Asterisks represent significance (*P*) of 2-tailed paired *t-*tests between Phase 1 and Phase 2: ** <.001

## DISCUSSION

Over the last decade, increased SED use and insufficient sleep in adolescents have been systematically reported and have been associated with various consequences on daytime performance and long-term difficulties (11, 13, 20, 38). However, to our knowledge, no study to date has investigated the impact of a restrictive exposure to SED screen in the evening on objective measures of sleep and wake performance in a large sample of adolescents. Here we tested whether a simple recommendation“no exposure to screens after 9 pm during two weeks”had beneficial consequences on sleep and performance in adolescents. Using subjective and objective measures of sleep on 183 adolescents (aged between 12 and 19 years old), we show that decreasing SED in the evenings preceding school days significantly advanced sleep onset and increased sleep duration, especially in older adolescents (14-19 years old). Secondly, we demonstrate that the Val158Met polymorphism of *COMT* gene may modulate these effects of reduced SED use after 9pm on sleep. Thirdly, we report improved daytime vigilance after reduced SED use. Together, these data provide unprecedented scientific evidence from a large population of adolescents that reducing SED use in the evening beneficially affects sleep and daytime vigilance.

### SED use in the evening delays sleep onset time and shortens sleep duration

Previous studies using questionnaires in children and adolescents have suggested a relationship between SED use in the evening and sleep habits (10, 38, 39). Here, using objective (i.e., actigraphy, melatonin profile) and subjective (i.e., daily diaries) measures, we confirm that time spent on SED after 9pm correlates with later sleep onset time, and consequently shorter total sleep duration, as wake up time does not change during school-days. Moreover, we provide further experimental support for this relation by demonstrating that decreasing SED use in the evening advances sleep onset time and increases sleep duration. Why does SED use affect sleep? SED use is time consuming, and would thus simply compete with sleep time, especially when wake-up time is constrained as this is the case for adolescents during school days (13). Yet, in our study, adolescents devoted a substantial amount of time doing off-screen activities, but only time spent on screen-based activities significantly correlated with sleep parameters, thus not corroborating the claim of SED use merely replacing sleep time (see also below). SED use also often implies activities that are known to increase stress and emotional arousal levels (e.g., social media, video games) (15, 40), which can impact bedtime hour as well as sleep initiation (41). Our data seem to suggest that indeed not all SED activities affected sleep to the same extent. Yet, while older teenagers spent more time on social media platforms, our SEM analysis revealed that social media use did not affect sleep duration more than other SED activities. Finally, SED may also influence sleep through the high spectral radiance blue-light emitted by the screen devices, which was shown to directly interfere with the circadian regulation by the suprachiasmatic nucleus via the retino-hypothalamic pathway (42). Exposure to SED thus delays the evening rise of the sleep-promoting hormone melatonin, leading to an increase in alertness and reduced sleepiness (17–19, 43, 44). Our results are in line with the latter hypothesis since all activities on SED (i.e., PC and smartphones) contributed significantly to the modulation of sleep duration, except watching TV. A sufficient distance between screen and eyes might indeed prevent sleep disruption due to the blue-light emitted by the screen (8). We could not detect a significant change in melatonin profile as a function of the intervention but note that this analysis was conducted on a very restricted number of participants (*N*=13) and should be considered as preliminary.

### Controlling SED use in the evening as an effective strategy to improve sleep and cognition in adolescents

In the present study, our sample of adolescents slept less (mean ± SEM; 7h33 ± 3 min during pre-school nights) than the 10 h (+/- 1 h) for school age (6-13 years) and 9 h (+/- 1 h) for teenager (14-17 years) recommended by the National Sleep Foundation (45), indicating a state of chronic sleep deprivation during the week (46), as further evidenced by a rebound of sleep (i.e., extended sleep duration) during weekend nights. Our data show that short sleep duration correlated with higher daytime fatigue, psychological distress and low mood rating (24). Moreover, chronic sleep deprivation at such a young age can put them at risk for the development of sleep and health disorders, such as depression, diabetes or obesity (21, 47, 48). For example, in our sample, both short sleep duration and extensive time on SED in the evening correlated with higher BMI. These alarming observations call for the development of strategies to extend sleep duration in adolescents. The efficient strategy would be to act on both bedtime and wake-up time, one being via changing school time (49), while the other would change pre-sleep behavior, SED use being a likely efficient target. In 2015, Harris et al. (50) reported that a restrictive use of SED after 10 pm in a sample of high school athletes did not improve sleep habits, mood, or physical and cognitive performance. As suggested by these researchers, it is plausible that preventing SED use after 10 pm was not effective because the tested population had early habitual bedtimes already. By contrast, in our study, we found that reducing SED use after 9 pm (i.e. following the recommendation) decreased sleep onset time, increased sleep duration and daytime vigilance. We also observed that the impact of the intervention on sleep was modulated by individual genetic background. Converging with our results on a general population of adolescents, a previous study showed that extending sleep duration by gradually advancing bedtime in teenagers exhibiting symptoms of chronic sleep reduction (*N*=28) led to earlier sleep onset and longer sleep duration, which had repercussions on cognitive performance (25). All age groups decreased their time spent using SED during Phase 2, with associated beneficial consequences on sleep for all groups except the youngest group (12-13 years old).

This observation might be explained by the low use of SED in the evening and earlier bedtime in younger participants during Phase 1, thus leaving less room for changes in SED use and the potential effects on sleep during Phase 2 (Fig. 3*B* and 3*D*). Parental supervision might therefore play a more significant role in this age group, thus limiting the impact of the intervention. Indeed, it has been shown that parental monitoring of bedtime during weeknights decreased with age and can be nearly absent for older teenagers (from 17 years old (51)). However, we did not measure parental control in our sample.

### Preliminary evidence for individual differences related to *COMT* activity

Our data suggest that the beneficial impact on sleep parameters might be genotype-dependent. Indeed, despite a similar decrease in SED use during Phase 2, the homozygote Met/Met *COMT* adolescents did not advance their sleep onset time, nor did they extend their sleep duration, compared to the other *COMT* genotype (Val/Val and Val/Met) individuals. *COMT* gene polymorphism has been associated with dopamine-related cognitive and emotional functioning and, more recently, to sleep-wake regulation (26, 33, 35). Met allele carriers have been shown to have lower *COMT* activity and thus higher dopaminergic signaling in prefrontal cortex compared to Val carriers (29, 52). Met carriers exhibited higher fast sleep spindle density during NREM (37) but also faster alpha-peak frequency and higher upper alpha band power during wakefulness as well as during REM and NREM (53, 54). Faster alpha-peak frequency and more activity in alpha band due to higher dopaminergic activity might explain why Met carrier have better performance in executive tasks than Val carrier (28, 55), and why they may be less vulnerable to the impact of sleep deprivation (35, 56). More generally, these findings suggest that the efficiency of such interventions will always be partly moderated by inter-individual differences, some of which might relate to genetic factors.

### Limitations

A first limitation of the present investigation and previous studies on SED use relates to the fact that SED activities were measured through questionnaires, scales or self-reports that rely on the participant’s own perception and willingness to communicate the information. It would thus be necessary in the future to find ethical and objective strategies to obtain these data, such as using applications that would measure all activities directly from the electronic devices. However, because our main results come from within-subjects comparisons between Phase 1 and Phase 2, the impact of such potential distortion in the data is limited. A second limitation concerns the relatively simple format of the diary on evening activities, which did not allow us to account for task switching or multitasking in the analyses. We thus cannot fully rule out that time spent on SED might be slightly over-estimated. A third limitation is the small number of participants for whom we could extract the melatonin profiles during Phase 2. This might have led to undetected differences between both phases and prevented the inspection of further links with other variables. These results should therefore be considered as preliminary. Lastly, we could not collect data about the psychosocial environment, economic status, or parental monitoring, all factors that may also modulate the impact of an intervention such as the one we used here.

Our findings have several implications. First, we confirmed with objective and subjective measures that technology-related behavior before sleep has a negative impact on sleep parameters. Second, we found that it is possible to extend sleep in adolescents by imposing a restriction on their SED use after 9 pm, with beneficial consequences on daytime vigilance. Finally, we suggest that COMT polymorphism may partly explain why some individuals benefit more (than others) from limited SED use. The present study represents a necessary step towards the development of strategies to improve chronic sleep deprivation related to SED use, which has recently emerged as a major health issue in adolescents.

## MATERIALS AND METHODS

### Protocol

The experimental design of the study included two successive two-week periods (**Fig. 1**), during which participants wore an actimeter and filled out daily questionnaires on sleep and time spent performing screen-based and off-screen activities. The first period (or Phase 1) served as a baseline assessment. During the second period (or Phase 2), participants were instructed to reduce their SED use after 9 pm and any change in the collected variables was examined. Experimenters met with the participants three times (or Visits), right before Phase 1, between the two phases, and after the completion of Phase 2. During Visit 1, the experimenters presented the experimental procedures and timeline to the participants; Visit 1 always took place in the participants’ school. Next, after two weeks of “normal” SED use, participants came to the lab for Visit 2, during which they attended an interactive workshop on sleep physiology, sleep disorders, and the importance of sleep on daytime functioning and health. Then, they filled out several questionnaires and performed the Sustained Attention to Response Task (SART). One salivary sample for genetic profiling was also collected. At the end of Visit 2, the instructions for the two following weeks (Phase 2) were explained to the participants. Specifically, they were asked to stop using SED after 9 pm on pre-school evenings, namely from Sunday to Thursday evenings. Finally, a brainstorming was conducted to help them come up with off-screen activities that they could engage in after 9 pm. Subjects who agreed to participate to Phase 2 committed symbolically (signed a declaration of participation) to follow the restrictive rule. Two weeks later, Visit 3 took place in the schools where all participants filled out several questionnaires and performed the SART. To characterize each participant’s melatonin profile and possible changes after Phase 2, salivary samples were collected at home during the night after Visits 1 and 3 (see Melatonin section below). This study was approved by the ethics review board of the Geneva University Hospitals. Adult participants - or the parents of participants under 18 years old - signed the informed consent before taking part in the study. All data collected were kept anonymous using personal identification codes.

### Participants

In total, 569 students between 12 and 19 years old (52.5 % girls; mean age ± SD: 15.35 ± 2.1), recruited from middle and high schools in Geneva (Switzerland), took part in at least Phase 1 of the study. To obtain reliable estimates of SED use and sleep habits, only participants who filled out all the questionnaires during Visit 2, wore the actimeter, and filled out daily diaries for at least 7 days were analyzed and were called “*Active*” participants for Phase 1. The same criteria were used for Phase 2, with the additional requirement that *Active* participants in Phase 2 also had to be *Active* in Phase 1. Participants who did not meet these criteria were called “*Passive*” and we only analyzed their data from the questionnaires and SART, i.e. data collected in the lab and in the classroom. Note that some participants did not participate to Visit 3, and are referred to as “*Drop-outs*”. Accordingly, there were 315 *Active* participants (64.4 % girls, mean age ± SD: 15.69 ± 2.12) and 254 *Passive* participants (37.7 % girls, mean age ± SD: 14.93 ± 2) for Phase 1. Out of the 315 *Active* participants from Phase 1, 183 (65.5 % girls; mean age ± SD: 15.74 ± 2.08) agreed to follow the restrictive rule and went on to Phase 2 of the study for at least 7 days. Thus, for Phase 2 (and comparisons between Phases 1 and 2), there were 183 *Active* participants, 284 *Passive* participants (43.3 % girls; mean age ± SD: 14.84 ± 1.94), and 102 *Drop-outs* (54.9 % girls; mean age ± SD: 16.1 ± 2.19).

To ensure that the results of the intervention were not influenced by a selection bias, *Active* subjects who participated to both phases (*N*=183) were compared to those who were only *Active* during Phase 1 (*N*=132). Age, gender breakdown, sleep parameters, and duration of evening activities during Phase 1 did not differ between these groups, suggesting that the data obtained during Phase 2 (and comparisons between Phases 1 and 2) were not confounded by a selection bias. To investigate the effect of Age on the relation SED use and sleep, we further separated participants into four age groups: 12-13, 14-15, 16-17 and 18-19 years old. Descriptive data (age, gender) about these four groups for Phase 1 and 2 can be found in **Table S2**.

### Measures

#### Sleep and media diaries

Every day, participants provided information about their sleep and their evening activities on SED and off-screen (either on paper version or via internet). For sleep, they reported their bedtime, time to fall asleep (sleep latency), wake-up time, out of bedtime, and the number of nocturnal awakenings. They also evaluated their sleep quality and morning mood using a 5-star rating system (from 1: very bad sleep to 5 stars: very good sleep, and from 1: very bad mood to 5 stars: very good mood). For evening activities, they reported the time spent (in minutes) on different screen-based activities and off-screen activities after 9 pm (from 9 pm until sleep onset). Screen-based activities were categorized as either social media (e.g., Facebook, WhatsApp, SMS, Snapchat), watching TV, watching videos (not on TV), playing games and computer activities (e.g., email, reading blogs, online homework). Off-screen activities comprised of time spent on homework (not using a computer), sports, and reading. We then calculated the average of each variable for pre-school nights versus weekend nights. “Pre-school nights” referred to the evenings and nights preceding school days (i.e., Sunday to Thursday). “Weekend nights” were the evenings and nights before weekend days (i.e., Friday and Saturday) where participants had no wake-up time constraint related to school.

#### Actigraphy

Participants wore an Actimeters GT3X+ (Actigraph, Pensacola FL, USA) on their non-dominant wrist non-stop for the two successive periods of two weeks (Phases 1 and 2). This device contains a triaxial accelerometer with a dynamic range of ± 8 G, a sampling rate of 30 Hz and data are stored in a raw non-filtered format in G’s directly into a non-volatile flash memory. Mean actigraphic data during 60 s epochs were scored as sleep or wake using an automatic detection algorithm (AARA; http://orbi.ulg.ac.be/handle/2268/173392). We, then, reviewed each night manually by comparing the sleep times indicated by the automatic detection with those reported by the participants in the diaries. In case of a mismatch greater than one hour for bedtime or out-of-bed time, the night was removed from further analysis. In case of a mismatch of less than one hour, bedtime hour was corrected starting at the first Sleep Bout detected after the subjective bedtime reported by the participant. Morning out-of-bed times were similarly considered, by either removing the night for mismatches greater than one hour, or by correcting it using the beginning of the first Wake bout after subjective out-of-bed time reported by the participant. From these data, we computed several sleep variables: sleep onset time (SO; first sleep bout longer than 5 minutes), total sleep period (TSP; period between SO and wake-up time) and sleep efficiency (SE in %: (TSP minus wake time intra-sleep)/total time in bed*100).

#### Questionnaires

All participants (*N*=569), whether they wore the actimeter, filled out the diaries, followed the restrictive rule or not, filled out two sets of questionnaires: one at the end of Phase 1 and one at the end of Phase 2. Besides questionnaires about their age, sex, height, weight, health status, consumption habits, evening activities habits, and academic performance, participants also answered the Chronic Sleep Reduction Questionnaire (CSRQ) (57), which contains questions about sleepiness, irritation, and loss of energy during the day. They also responded to the Kessler Psychological Distress Scale (K6) (58) that quantifies non-specific psychological distress including anxiety, depression, despair.

#### Sustained Attention to Response Task (SART)

Participants performed the SART on tablet computers at the end of each phase. Due to technical problems with tablet early in the study, we included data from 454 (out of 569) participants in the final analyses. The SART consists of a GO/NO GO task, which measures sustained attention and vigilance performance (59). Participants were shown digits from 0 to 9 presented one at a time, each for 250 ms, and they had to tap the screen as quickly as possible when a digit appeared (GO) except if it was the digit “3” (NO GO). A fixation cross was presented for 900 ms after each digit. A total of 250 single digits were presented in a pseudo-random order excluding the immediate succession of the same number, with every digit appearing 25 times. The duration of the task was 4.3 min. We analyzed the number of commission errors (GO when the “3” appeared), the number of omissions (not responding to a “GO” signal), and the sum of both types of errors (i.e., total number of errors) (60). We calculated mean reaction times (RTs) for corrects responses, but also the 75^th^ percentile (slowest) of the distribution of the RTs, which has been shown to best reflect deterioration of psychomotor vigilance and wake instability due to augmented sleep pressure (22, 61). Indeed, the slowest RTs have been shown to be more sensitive to sleep deprivation than faster RTs (62).

#### Melatonin

Five saliva samples were collected twice in order to assess individual melatonin profiles before Phase 1 and after Phase 2 (Fig. 1). Collection was performed at home by the participants using the SalivaBio Oral Swab (SOS; Salimetrics; Suffolk, UK) method. They were asked to collect saliva every hour, starting 4 h before their usual bedtime and finishing with the last one collected 1 h after their usual bedtime. They were instructed to avoid eating one hour before the first extraction and between the five saliva samples. They were also asked to avoid drinking alcohol and energy drinks after 3 pm, and also to avoid too much sweet (e.g. chocolate, banana) or sour (e.g. lemon) food during the last meal. They were asked to place the tubes with the saliva samples in their fridge and to bring them at their school the next morning, where they were collected by one experimenter. Salivary samples were centrifuged briefly to collect supernatant at the bottom of the tube (only when there was at least four samples for one evening) and the pre-processed samples were then kept at minus 80 ^°^C until assays were performed. The quantitative determination of melatonin in saliva was then obtained using enzyme-linked immunosorbent assay (ELISA) kits (Direct Saliva Melatonin ELISA; Bühlmann Laboratories, Allschwil, Switzerland). Hour of Dim Light Melatonin Onset (HDLMO) was calculated using the hockey-stick method (63). We were able to successfully analyze melatonin profile for 70 *Active* participants for Phase 1, while only 13 *Active* participants had melatonin profiles for both Phase 1 and 2.

#### Genotyping

Saliva samples were collected during Visit 2 and consisted in spitting at least 3 ml of saliva in a tube. Tubes were kept at minus 80 ^°^C until analyses were performed. Total genomic DNA was extracted from saliva samples using the Gentra-Puregene kit (QIAGEN; Venlo, Netherlands). We genotyped the A/G SNP of *COMT* gene (ID of the National Center for Biotechnology Information dbSNP database: rs4680, location: chr 22: 19951271) using the TaqMan platform (https://ige3.genomics.unige.ch) for allelic discrimination (Applied Biosystems). PCR amplification was performed on 384-well plates using TaqMan Predesigned SNP Genotyping Assays (Applied Biosystems) and conditions recommended by the manufacturer. Reactions were analyzed by individuals blinded to subject using the Applied Biosystems TaqMan 7900HT system and the sequence detection system software v2.2.1. All samples were genotyped in triplicate, with 100% concordance. A total of 203 samples were successfully analyzed for *COMT*.

#### Statistical Analyses

Statistical analyses were performed using IBM SPSS Statistics for Windows (Version 23.0. Armonk, NY: IBM Corp).Using multivariate linear regressions, we examined associations between time spent on screen-based activities or off-screen activities after 9pm and several sleep parameters. Repeated measures analyses of variance (ANOVA) with post-hoc pairwise t-tests were used to specify main and interaction effects due to the instruction. Thus, most ANOVAs included a repeated measure factor Phase (Phase 1, Phase 2) as within-subject factor. In order to test for any effect of age or *COMT* gene, some ANOVAs were computed with between-subjects factor Age Group (12-13, 14-15, 16-17, 18-19 years old) or *COMT* Gene (Val/Val, Val/Met, Met/Met polymorphisms). Degrees of freedom were corrected according to Greenhouse-Geisser, when appropriate. The level of significance was set to a p-value < 0.05. To further examine the impact of the different types of screen-based activities performed after 9 pm on sleep duration during Phase 1, we used a Structural Equation Modeling (SEM) approach (multivariate path analysis in SPSS AMOS Version 23.0. Armonk, NY: IBM Corp).

## ACKNOWLEDGMENT

We thank Dr. Swann Pichon, Laurence Schenkel Coste, Soledad Valera-Kummer, Philippe Lavorel and Dr. Dagmar Haller-Hester for their valuable inputs at the beginning of the project. This work was supported by the Gertrude von Meissner Foundation and by the National Center of Competence in Research (NCCR) Affective Sciences financed by the Swiss National Science Foundation (grant number: 51NF40-104897) and hosted by the University of Geneva.

