## Supplementary Materials for "Imposing a curfew on the use of screen electronic devices improves sleep and daytime vigilance in adolescents"

SUPPLEMENTAL INFORMATION

Table S1: COMT genotypes repartition (*N*=121) and characteristics (gender, age, body mass index, Phase 1 parameters).

| Participants (*N*=121) with genetic (*COMT*) profiling | | | |
|---|---|---|---|
| *Allele* | **Val/Val** | **Val/Met** | **Met/Met** |
| *N* | 37 | 61 | 23 |
| *Girls / Boys* | 22 / 15 | 43 / 18 | 16 / 7 |
| *Age (mean ± SD)* | 16.13 ± 2 | 15.54 ± 2.1 | 15.69 ± 1.52 |
| *BMI (mean ± SD)* | 20.88 ± 2.85 | 19.83 ± 2.53 | 20.33 ± 3.11 |
| Phase 1 parameters (mean ± sem) | | | |
| *-SED use after 9 pm (min)* | 86 ± 8 | 70 ± 6 | 88 ± 12 |
| *-Offscreen activities after 9 pm (min)* | 53 ± 6 | 52 ± 5 | 59 ± 9 |
| *-Sleep duration (hours)* | 7h25 ± 0.13 | 7h34 ± 0.1 | 7h28 ± 0.13 |
| *-Sleep efficiency (%)* | 89.1 ± 0.7 | 89.1 ± 0.6 | 89.8 ± 0.9 |
| *-Daily mood (from 1 to 5 scale)* | 3.58 ± 0.09 | 3.49 ± 0.07 | 3.51 ± 0.11 |

There was no significant difference between the three groups during Phase 1.

Table S2: Repartition between *Active* and *Passive* participants and their characteristics (gender, age)

| Participants included in PHASE 1 | | | | | |
|---|---|---|---|---|---|
| Total Participants | Total | 12-13 y.o | 14-15 y.o | 16-17 y.o | 18-19 y.o |
| N | 569 | 140 | 168 | 138 | 123 |
| Girls \| Boys | 299 \| 270 | 72 \| 68 | 70 \| 98 | 83 \| 55 | 74 \| 49 |
| Age (mean ± SD) | 15,35 ± 2,1 | | | | |
| Active Participants | Total | 12-13 y.o | 14-15 y.o | 16-17 y.o | 18-19 y.o |
| N | 315 | 68 | 72 | 93 | 82 |
| Girls \| Boys | 203 \| 112 | 42 \| 26 | 42 \| 30 | 62 \| 31 | 57 \| 25 |
| Age (mean ± SD) | 15,69 ± 2,12 | | | | |
| Passive Participants | Total | 12-13 y.o | 14-15 y.o | 16-17 y.o | 18-19 y.o |
| N | 254 | 72 | 96 | 45 | 41 |
| Girls \| Boys | 96 \| 158 | 30 \| 42 | 28 \| 68 | 21 \| 24 | 17 \| 24 |
| Age (mean ± SD) | 14,93 ± 2 | | | | |

| Participants included in both PHASES 1 & 2 | | | | | |
|---|---|---|---|---|---|
| Total Participants | Total | 12-13 y.o | 14-15 y.o | 16-17 y.o | 18-19 y.o |
| N | 467 | 120 | 151 | 111 | 85 |
| Girls \| Boys | 243 \| 224 | 61 \| 59 | 64 \| 87 | 67 \| 44 | 51 \| 34 |
| Age (mean ± SD) | 15,02 ± 1,9 | | | | |
| Active Participants | Total | 12-13 y.o | 14-15 y.o | 16-17 y.o | 18-19 y.o |
| N | 183 | 36 | 43 | 58 | 46 |
| Girls \| Boys | 120 \| 63 | 17 \| 19 | 28 \| 15 | 43 \| 15 | 32 \| 14 |
| Age (mean ± SD) | 15,73 ± 2 | | | | |
| Passive Participants | Total | 12-13 y.o | 14-15 y.o | 16-17 y.o | 18-19 y.o |
| N | 284 | 84 | 108 | 53 | 39 |
| Girls \| Boys | 123 \| 161 | 44 \| 40 | 36 \| 72 | 24 \| 29 | 19 \| 20 |
| Age (mean ± SD) | 14,84 ± 1,95 | | | | |

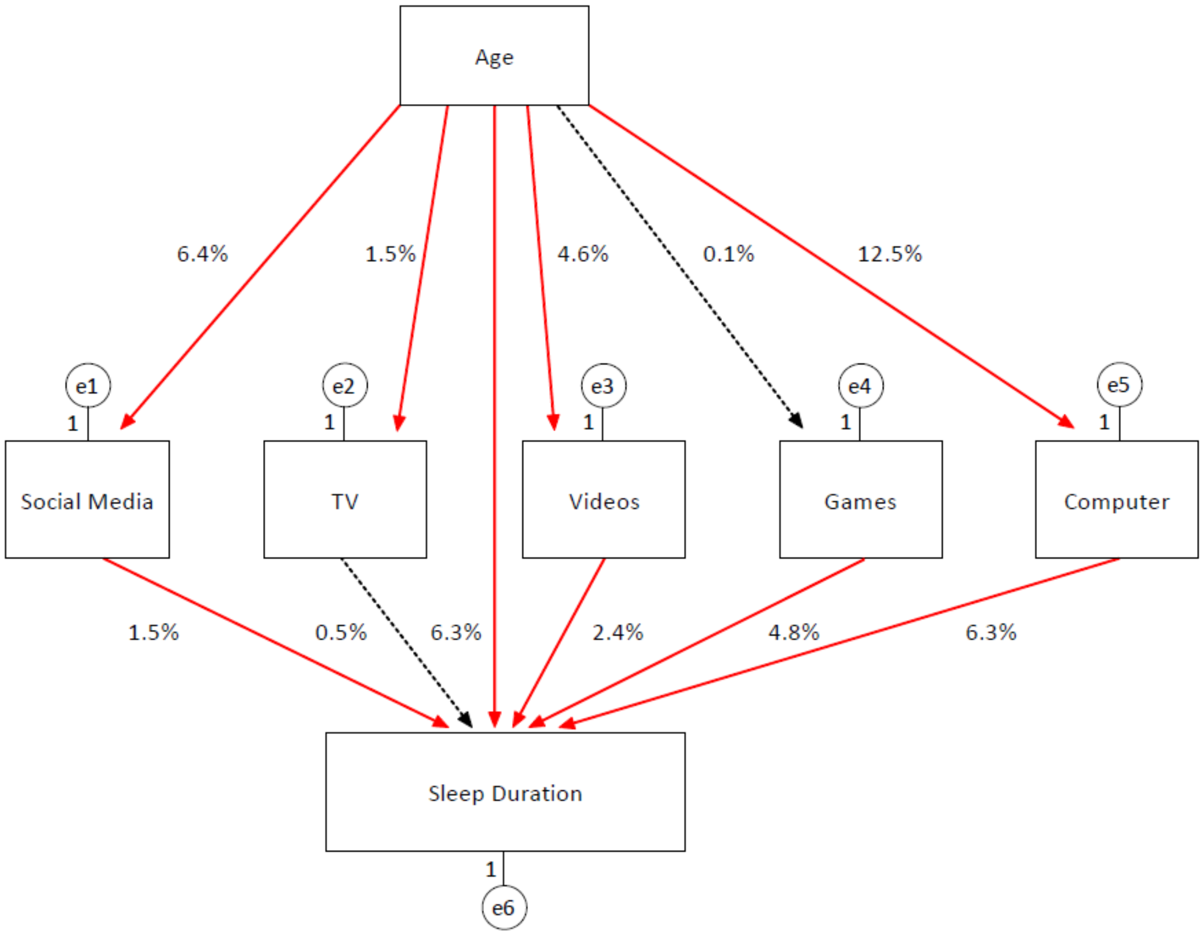


**Fig. S1: Contribution of age and type of SED activities after 9 pm to sleep duration during pre-school nights using Structural Equation Modeling**

Age and time spent on each SED activity after 9 pm collectively explained 37.8% of the variance of the variable sleep duration. While age and time spent on computer, games, videos and social media after 9 pm had significant unique contributions to sleep duration (red arrows), time spent watching TV after 9 pm did not significantly affect sleep duration (dotted black arrow).
